# Bifunctional small molecules that induce nuclear localization and targeted transcriptional regulation

**DOI:** 10.1101/2023.07.07.548101

**Authors:** William J. Gibson, Ananthan Sadagopan, Veronika M. Shoba, Amit Choudhary, Matthew Meyerson, Stuart L. Schreiber

**Author notes:** These authors contributed equally (William J. Gibson, Ananthan Sadagopan). These authors contributed equally and jointly supervised this work (Matthew Meyerson, Stuart L. Schreiber). Correspondence should be addressed to W.J.G., M.M., and S.L.S.

## Abstract

The aberrant localization of proteins in cells is a key factor in the development of various diseases, including cancer and neurodegenerative disease. To better understand and potentially manipulate protein localization for therapeutic purposes, we engineered bifunctional compounds that bind to proteins in separate cellular compartments. We show these compounds induce nuclear import of cytosolic cargoes, using nuclear-localized BRD4 as a “carrier” for co-import and nuclear trapping of cytosolic proteins. We use this system to calculate kinetic constants for passive diffusion across the nuclear pore and demonstrate single-cell heterogeneity in response to these bifunctional molecules, with cells requiring high carrier to cargo expression for complete import. We also observe incorporation of cargoes into BRD4-containing condensates. Proteins shown to be substrates for nuclear transport include oncogenic mutant nucleophosmin (NPM1c) and mutant PI3K catalytic subunit alpha (PIK3CA_E545K_), suggesting potential applications to cancer treatment. In addition, we demonstrate that chemical-induced localization of BRD4 to cytosolic-localized DNA-binding proteins, namely, IRF1 with a nuclear export signal, induces target gene expression. These results suggest that induced localization of proteins with bifunctional molecules enables the rewiring of cell circuitry with significant implications for disease therapy.

## Introduction

Cells partition their components into separate compartments to regulate information flow and to concentrate biomolecules. Eukaryotic cells possess two main compartments, the cytosol, and the nucleus. The trafficking of proteins across the nuclear membrane can be either passive, in the case of proteins less than 90 kDa, or facilitated by active transport^1,2^. Recent research further emphasizes the importance of protein spatial localization, as demonstrated by the discovery and diverse functions of phase-separated biomolecular condensates^3^. The translocation of proteins from one compartment to another is the defining event in several biological cascades. Examples include the release of cytochrome C from the mitochondrial intermembrane space to the cytosol to initiate apoptosis, or conversely the sequestration of eIF2α to membraneless stress granules for translational pausing, facilitating survival upon cellular stress^4,5^. Nuclear transport in particular is often a potent signaling event; many transcription factors typically reside in the cytosol but translocate to the nucleus upon activation. These transcription factors include the FoxO, STAT, and nuclear hormone receptor families of proteins^6–8^.

Given the diverse biological processes that are impacted by alternative protein localization, we sought to develop molecules that can control protein localization. We hypothesized that chemical-induced proximity could be used to induce the co-localization of proteins that normally exist in opposing compartments and thereby translocate one from its normal location to a new one. Earlier work has demonstrated that altering the localization of proteins is possible using the FKBP/FRB system^9^. We aimed to explore whether bifunctional compounds could be used to move proteins without the use of a dual-tagging system by leveraging a small-molecule binder of the endogenous protein BRD4.

We reveal bifunctional small molecules capable of inducing protein translocation from the cytosol to the nucleus. Such molecules may have important applications in human health, including cancer treatment.

## Results

### Chemical-induced proximity with BRD4 rapidly induces nuclear localization of the targeted protein

We hypothesized that chemical-induced proximity of a protein of interest with a constitutively nuclear protein would induce nuclear localization of the target protein (Fig. 1A). To achieve this, we sought out proteins that are abundant, nuclear localized and have well-validated small-molecule binders that could be functionalized. Bromodomain-containing protein 4 (BRD4) is one such protein, and (+)-JQ1, a small-molecule binder of BET-containing proteins including BRD4, has been used to make a variety of bifunctional compounds^10^. We synthesized a bifunctional molecule consisting of JQ1 and AP1867, a high affinity (K_d_ = 94 pM) ligand of FKBP^F36V^,^11^ with a PEG_2_-diamine linker: <u>N</u>uclear Import and <u>C</u>ontrol of <u>E</u>xpression compound 1 (**NICE-01**, Fig. 1B).

**Figure 1:**
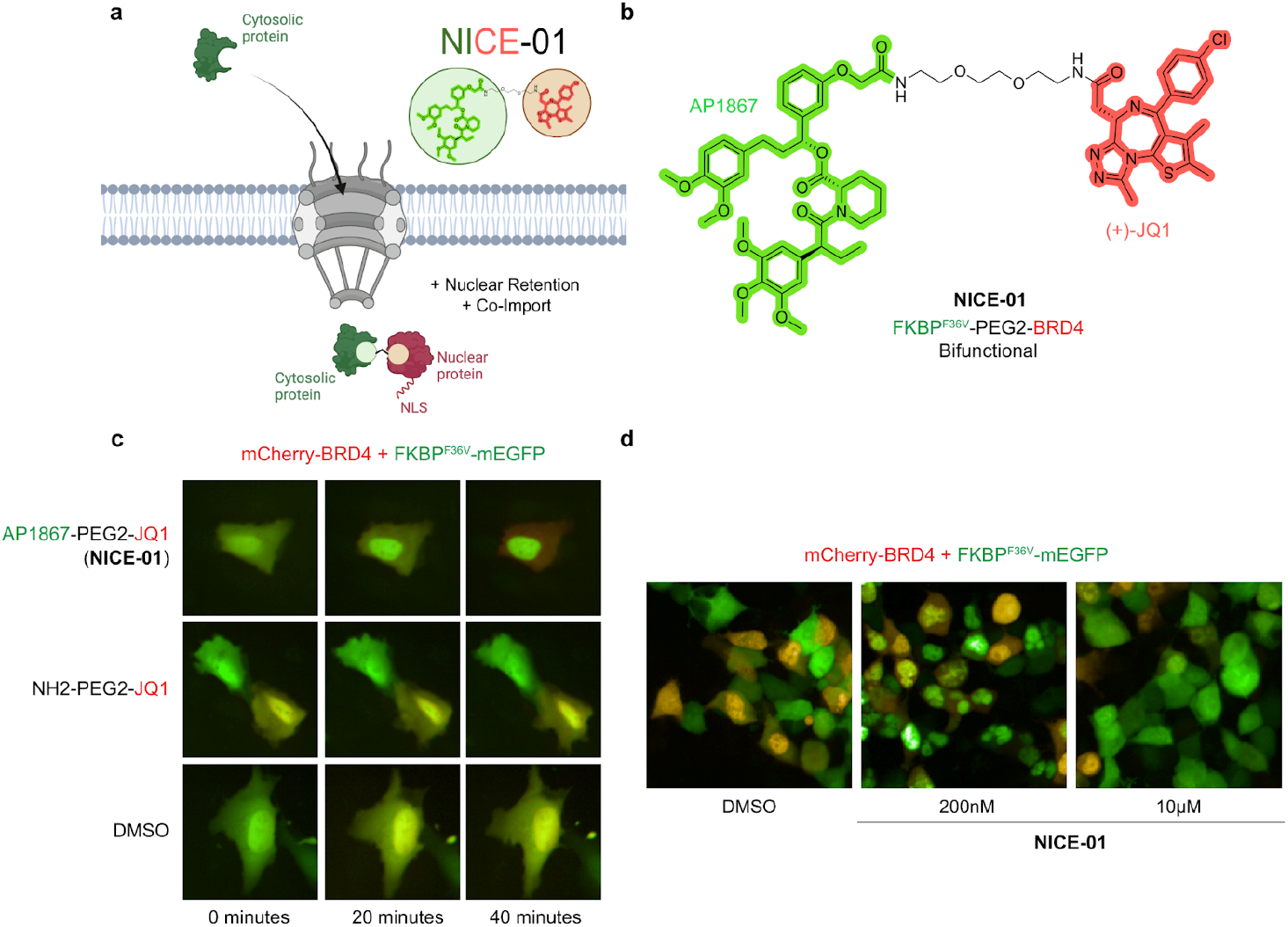
Rapid import of GFP into the nucleus by AP1867-PEG2-JQ1 (NICE-01). (A) Schematic of the <u>N</u>uclear Import and <u>C</u>ontrol of <u>E</u>xpression (NICE) concept. A bifunctional molecule allows import of a cytosolic protein of interest into the nucleus. (B) Structure of AP1867-PEG2-JQ1. (C) U2OS cells co-transfected with mCherry-BRD4 and FKBP^F36V^-mEGFP, treated with **NICE-01** (200 nM), NH2-PEG2-JQ1 (200 nM), or DMSO, and imaged for 40 minutes. (D) 293T cells co-transfected with mCherry-BRD4 and FKBP^F36V^-mEGFP, treated with DMSO or **NICE-01** (200 nM, 10 μM) and imaged 1-hour after addition.

Cells co-transfected with FKBP^F36V^-mEGFP and mCherry-BRD4 showed rapid translocation of FKBP^F36V^-EGFP to the nucleus following addition of **NICE-01** (200 nM, 40 minutes; Fig. 1C). One defining characteristic of bifunctional compounds, as opposed to molecular glues, is the observation of a dose-response that decreases at very high compound concentrations called the “hook effect”^12^. To confirm that the observed translocation is due to the bifunctional nature of **NICE-01**, we performed a dose-titration and found that nuclear translocation does not occur at a higher compound concentration of 10μM (Fig. 1D).

Recent studies have highlighted the importance of stoichiometry of the proteins involved in ternary complex formation for the biological activity of bifunctional molecules^13^. We also find that the stoichiometry of the components of the ternary complex was key to nuclear import. Indeed, we observed weaker FKBP^F36V^-mEGFP import when reliant on endogenous BET-containing proteins, with lack of mEGFP cytosolic exclusion (Fig. S1), likely related to the exceptionally high protein levels achieved through transient transfection compared to endogenous proteins. This observation suggests that cell specificity of nuclear import is quantitatively influenced by expression of the nuclear-localized carrier. Ligand affinity appears important as well. Bifunctional molecules constructed from synthetic ligand of FKBP (SLF), which binds with low micromolar affinity to wild-type FKBP^14^, instead of AP1867, required much higher concentrations (5 μM) to induce nuclear localization of FKBP-mEGFP in the presence of mCherry-BRD4 (data not shown).

mEGFP nuclear import may conceivably occur through two mechanisms. First, the protein of interest may co-import with BRD4 in the presence of bifunctional molecules, like how smaller Mediator complex subunits without NLSs “piggyback” off of larger NLS-containing subunits^15^. Second, the protein of interest may passively diffuse across the nuclear pore; in this case, the bifunctional molecule shifts the equilibrium towards the nucleus by sequestering the nuclear fraction with BRD4 thereby preventing export (“nuclear trapping”). To distinguish between these possibilities, we tested the cancer-driver PIK3CA^E545K^ mutant in our system, which is too large to passively diffuse across the nuclear pore (110 kDa; 149 kDa with FKBP^F36V^-mEGFP N-terminal fusion).^2^ **NICE-01** was able to import FKBP^F36V^-mEGFP-PIK3CA^E545K^ in cells co-transfected with mCherry-BRD4 (Fig. S2), indicating that co-import occurs. However, the time required for import (~3 hours) was much longer than FKBP^F36V^-mEGFP, consistent with nuclear trapping predominating on shorter timescales. While nearly all cells co-transfected with FKBP^F36V^-EGFP and mCherry-BRD4 showed complete nuclear localization of GFP within 1 hour, only about half of cells transfected with FKBP^F36V^-mEGFP-PIK3CA^E545K^ showed nuclear localization within 3 hours, and often only partial nuclear localization at steady state. This observation demonstrates that not all proteins will be equally amenable to nuclear import.

As diffusion of protein cargos across the nuclear membrane is required for nuclear trapping on short time scales, we hypothesized that the kinetic parameters of protein diffusion across the nuclear membrane could be calculated using the observed shift in FKBP^F36V^-mEGFP concentration upon compound treatment. We constructed a model of the steps in nuclear relocalization (Fig. S3; Supplementary Information). The fraction of cytosolic FKBP^F36V^-mEGFP was quantified for individual cells. Nuclear import showed first-order kinetics with respect to FKBP^F36V^-mEGFP fraction in the cytoplasm after a short initial period (during which nuclear FKBP^F36V^-mEGFP forms a ternary complex with mCherry-BRD4 and **NICE-01**); *k*_import_ was estimated to be 0.0266 min^-1^ (Fig. S3). Therefore, half of all cytosolic FKBP^F36V^-mEGFP (39.4 kDa) crosses the nuclear membrane every 26 minutes in the presence of **NICE-01**. This estimate is near the rate of passive import of similarly sized maltose-binding protein (MBP, 42.5 kDa) in HeLa cells (*k*_import_ = 0.0132 min^-1^, t_1/2_ = 53 minutes)^16^. These data indicate that a bifunctional compound that binds BRD4 induces nuclear import of target proteins.

### Nuclear localizing bifunctional compounds correct the localization of the cancer-associated nucleophosmin protein and alter nuclear condensates

The most common subtype of acute myeloid leukemia is defined by heterozygous mutations in the NPM1 gene that cause its aberrant cytosolic localization (NPM1c)^17^. Mutations in NPM1 occur exclusively at the C-terminus of the protein and induce a frameshift that simultaneously abolishes a nucleolar localization signal and introduces a neo-nuclear export signal (NES). The neo-NES causes exportin-1 (XPO1, CRM1) to bind and export the protein from the nucleus, shifting the equilibrium of protein localization to a predominantly cytosolic pattern in a dominant negative manner as NPM1c oligomerizes with wild-type NPM1^18–20^. Prior work has suggested that re-localizing the cytoplasmic mutant NPM1c into the nucleus can reduce the expression of HOX genes, thereby inducing leukemic differentiation and cell death^21^. We sought to determine whether nuclear relocalizing bifunctional compounds could be used to relocalize this mutant NPM1c into the nucleus. Cells co-transfected with FKBP^F36V^-mEGFP-NPM1c and mCherry-BRD4 showed rapid re-localization of FKBP^F36V^-mEGFP-NPM1c into the nucleus within minutes upon treatment with **NICE-01** (Fig. 2A-B, *P* = 4.4e-185 for Mann-Whitney U test between distribution cellular mEGFP-mCherry correlation before compound addition and 25 minutes after **NICE-01** addition).

**Figure 2:**
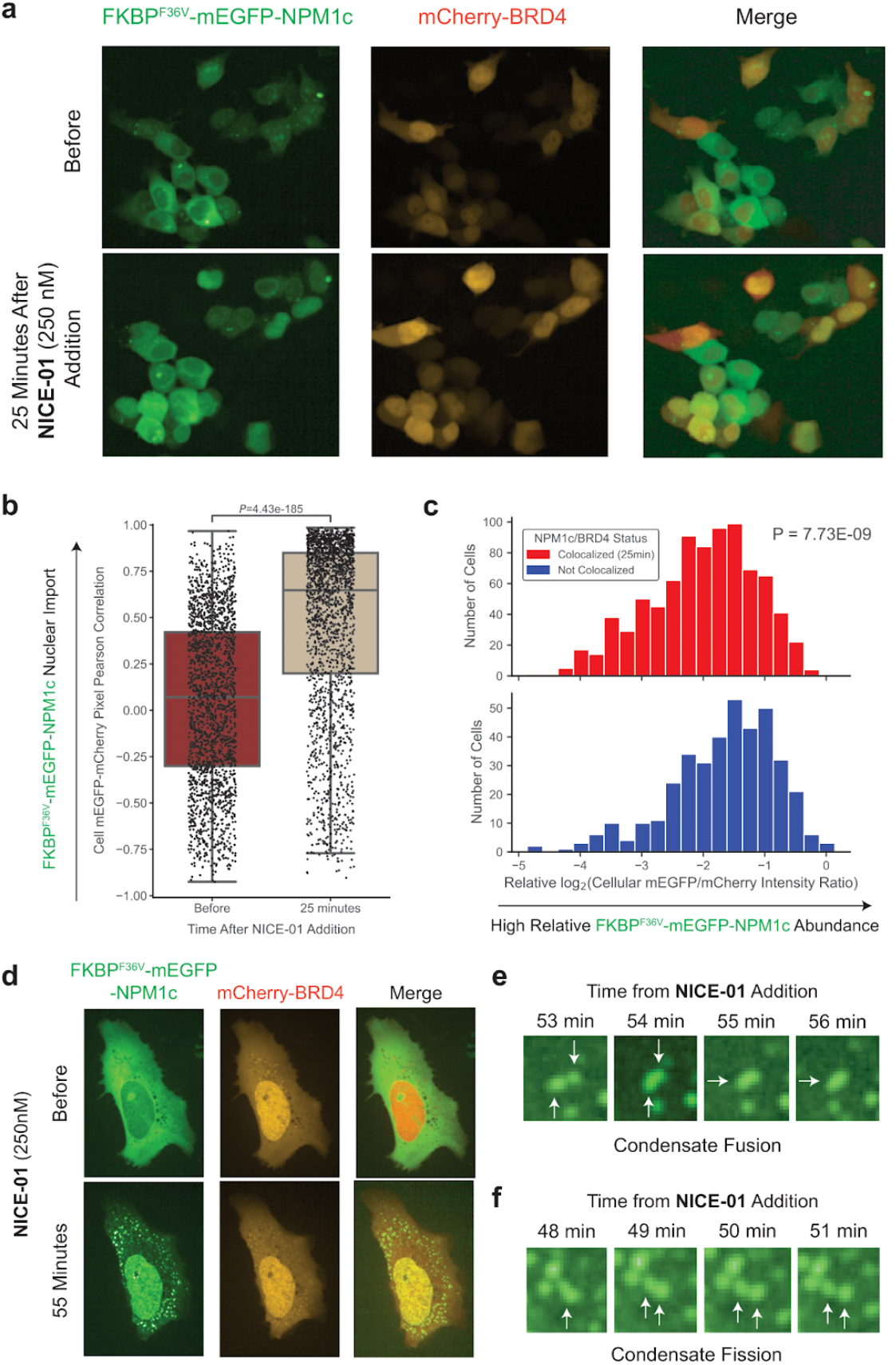
Import of cytosolically localized NPM1c into nuclear BRD4 condensates upon addition of bifunctional molecule NICE-01. (A) 293T cells co-transfected with FKBP^F36V^-EGFP-NPM1c and mCherry-BRD4 before (top) and 25 minutes after (bottom) **NICE-01** treatment (250 nM). (B) Boxplot of cellular mEGFP-mCherry pixel intensity correlation, before and 25 minutes after **NICE-01** treatment (250 nM, *P*-value calculated from Mann-Whitney U test). Each dot represents one cell. (C) Histograms of relative log_2_(integrated mEGFP cellular intensity / integrated mCherry cellular intensity) for cells with colocalized mEGFP and mCherry (mEGFP-mCherry pixel intensity correlation coefficient: R ≥ 0.75) or non-colocalized (mEGFP-mCherry pixel intensity correlation coefficient: R < 0). *P*-value calculated from Kolmogorov-Smirnov test. (D) High resolution images of U2OS cells co-transfected with FKBP^F36V^-EGFP-NPM1c and mCherry-BRD4 before (top) and 55 minutes after (bottom) **NICE-01** treatment (250 nM). (E) Image of two condensates fusing from panel (**D**), (F) Image of condensate undergoing fission from panel (**D**).

Interestingly, not all cells showed a uniform response to treatment with compound; duration required for mEGFP nuclear import varied, and nuclear relocalization was not observed in all cells. We observed that cells with low degree of NPM1c import often had lower mCherry-BRD4 intensity. We hypothesized that an excess of BRD4 might be required for efficient nuclear import. At an intermediate time point (25 minutes), we identified a significantly higher ratio of FKBP^F36V^-EGFP-NPM1c to mCherry-BRD4 in cells that did not respond (Fig. 2C, *P* = 7.7e-9 for Kolmogorov-Smirnov test between distribution of log_2_(cellular mEGFP/mCherry ratio) for mEGFP-mCherry colocalized versus mEGFP-mCherry non-colocalized cells). Taken together, these data suggest that the stoichiometry of components of the ternary complex can impact the phenotypic response to bifunctional compounds and may explain observed single-cell phenotypic heterogeneity.

We hypothesized that bifunctional compounds could enforce co-localization of proteins not only within macro-compartments, but also within smaller intracellular compartments such as phase-separated condensates. We performed high-resolution microscopy imaging every minute in U2OS cells co-transfected with NPM1c and BRD4, treated with **NICE-01**, and confirmed that NPM1c in the cytosol and in nucleolar condensates moved into BRD4 condensates, mostly in the nucleus and sometimes in the cytosol (Fig. 2D; Supplementary Movie 1). We confirmed these structures were condensates by observing them undergo fusion and fission over several minutes (Fig. 2E-F). These findings indicate proximity-inducing molecules can alter condensate composition in a targeted manner.

We next sought to determine if other nuclear carriers could be used to induce the localization of NPM1c into the nucleus and we hypothesized that p53 could be one such carrier. 293T cells were transfected with FKBP^F36V^-mEGFP-NPM1c and Halo-p53^R273H^-mCherry, which to our surprise demonstrated that Halo-p53^R273H^-mCherry was localized to the cytoplasm only when co-transfected with FKBP^F36V^-mEGFP-NPM1c (Fig. 3A). These findings are corroborated by a recent report^22^.

**Figure 3:**
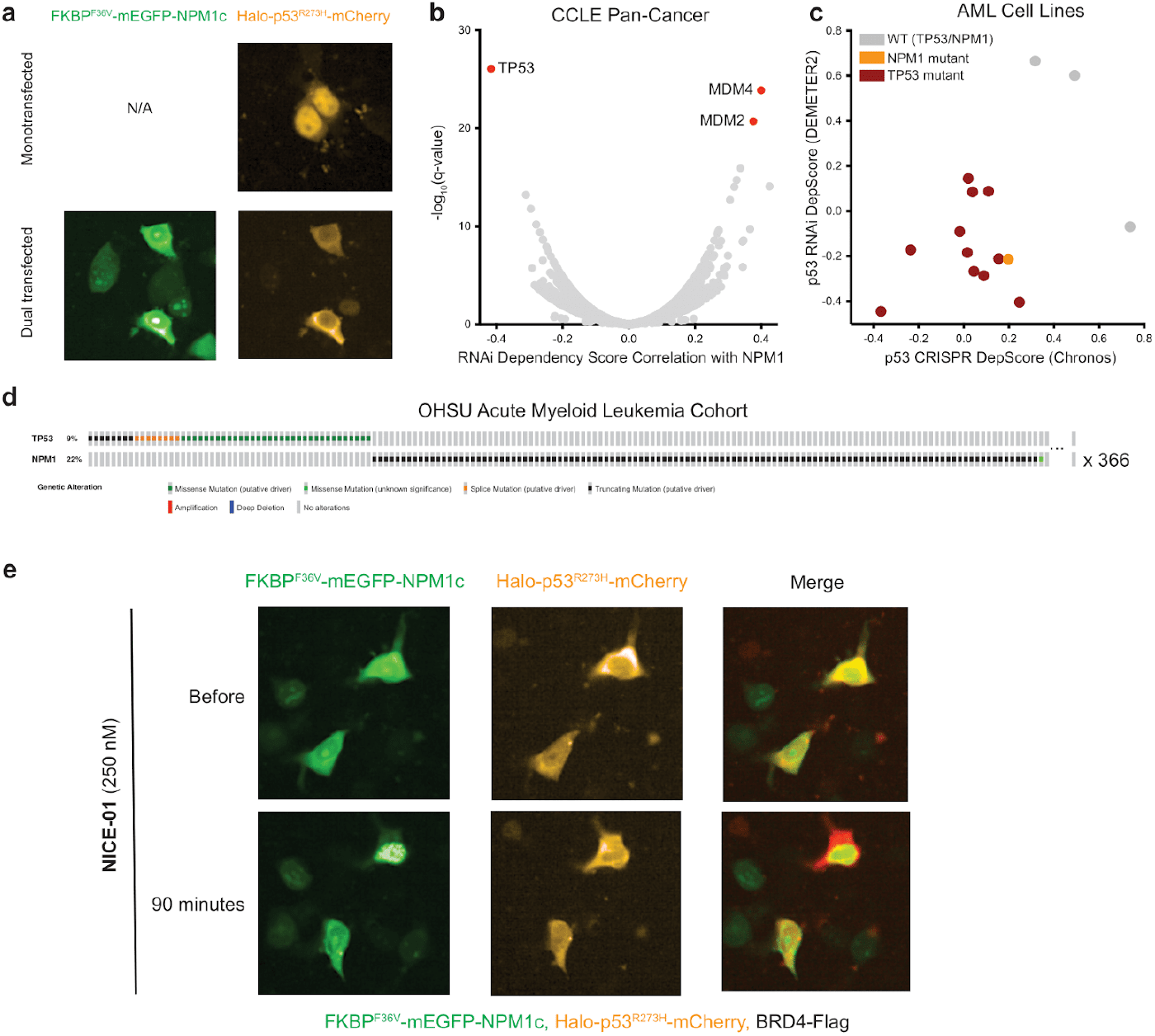
NPM1c exports p53 out of the nucleus and fails to import p53 upon addition of bifunctional compound. (A) 293T cells imaged following transfection with Halo-p53^R273H^-mCherry or co-transfection with FKBP^F36V^-mEGFP-NPM1c and Halo-p53^R273H^-mCherry. (B) RNAi co-dependency analysis of NPM1 across DepMap. (C) CRISPR and RNAi TP53 dependency score in AML cell lines across DepMap, colored by *TP53*/*NPM1* mutation status. (D) *NPM1* and *TP53* comut plot across OHSU acute myeloid leukemia cohort from cBioPortal. (E) 293T cells co-transfected with FKBP^F36V^-mEGFP-NPM1c, BRD4-Flag, and Halo-p53^R273H^-mCherry, treated with **NICE-01** (250 nM) and imaged for 90 minutes. Top right cell displays FKBP^F36V^-mEGFP-NPM1c import but lack of p53 co-import.

Cytosolic p53 is physically separated from DNA, likely inhibiting much of its tumor suppressive activity. Therefore, NPM1c-mediated p53 export could represent a significant contribution to the oncogenicity of the mutation, which is not well understood. This is supported by three lines of evidence. First, the genes for p53 and its E3 ligases, Mdm2 and Mdm4, are the top co-dependencies of *NPM1* inactivation across the Cancer Cell Line Encyclopedia^23^ (RNAi Achilles + DRIVE + Marcotte, DEMETER2; Fig. 3B). Second, the only NPM1c-mutant AML cell line in the Cancer Cell Line Encyclopedia, OCI-AML3, which has wild-type *TP53*, displays the smallest fitness gain upon *TP53* knockout of all *TP53* wild-type AML cell lines (Fig. 3C). Third, NPM1c mutations are mutually exclusive with p53 mutations in AML (*P* = 1.2e-6 for Fisher exact test on OHSU AML cohort, Fig. 3D)^24^. However, **NICE-01** could not relocalize

Halo-p53^R273H^-mCherry in 293T cells expressing BRD4-Flag and FKBP^F36V^-mEGFP-NPM1c, suggesting that p53 is not consistently physically tethered to NPM1c in such a way that it must always co-localize (Fig. 3E).

### Chemical-induced import of cytosolic transcription factor and induction of transcriptional response

The nuclear membrane sometimes acts as a barrier to prevent the activation of transcription factors that normally reside in the cytosol^25^. Upon activation, these transcription factors translocate to the nucleus where they can bind their target sites and induce transcription. Transcription factors that follow this pattern include nuclear hormone receptors ER, AR, and some of the STAT and IRF family proteins^7,8,26^. We hypothesized that **NICE-01** could induce the nuclear import and transcriptional activation of such cytosolic transcription factors. We selected the prototypical IRF family member, IRF1, for its restricted and well-described transcriptional profile^27,28^. An NES was appended to IRF1 to aid in the detection of nuclear translocation and facilitate the use of this construct in an “up-assay”.

In the absence of bifunctional compound, FKBP^F36V^-mEGFP-IRF1-NES was excluded from the nucleus (Fig. S4). We observed rapid mEGFP import (<20 minutes) in cells co-expressing mCherry-BRD4 and FKBP^F36V^-mEGFP-IRF1-NES when treated with **NICE-01** (250 nM). However, we could not detect FKBP^F36V^-mEGFP-IRF1-NES import in the absence of exogenous BRD4 by light microscopy, likely due to a large excess of FKBP^F36V^-mEGFP-IRF1-NES compared to BRD4 (Fig. S4). Though not visibly apparent, we hypothesized that nuclear residence time of transcriptionally active FKBP^F36V^-mEGFP-IRF1-NES was substantially increasing under these conditions and FKBP^F36V^-mEGFP-IRF1-NES could be getting incorporated into BRD4 condensates driving expression of IRF1 target genes. Consistent with this hypothesis, we found significant upregulation of IRF1 ChIP-seq targets^29^ including *IFI6, MT2A, IFI1*, and *B2M* by RNA-seq (*P*=3.4e-9 for one sample t-test versus μ = 0) in 293T cells treated with **NICE-01** (250 nM) and expressing FKBP^F36V^-mEGFP-IRF1-NES (Fig. 4A). We also observed significant positive enrichment of the hallmark interferon response gene sets by gene set enrichment analysis^30^ (IFNα, NES=3.31, q<0.001; IFNγ, NES=3.26, q<0.001; Fig. 4B). Therefore, we demonstrate chemical-induced localization of BRD4 to cytosolic transcription factors is sufficient to induce expression of their target genes.

**Figure 4:**
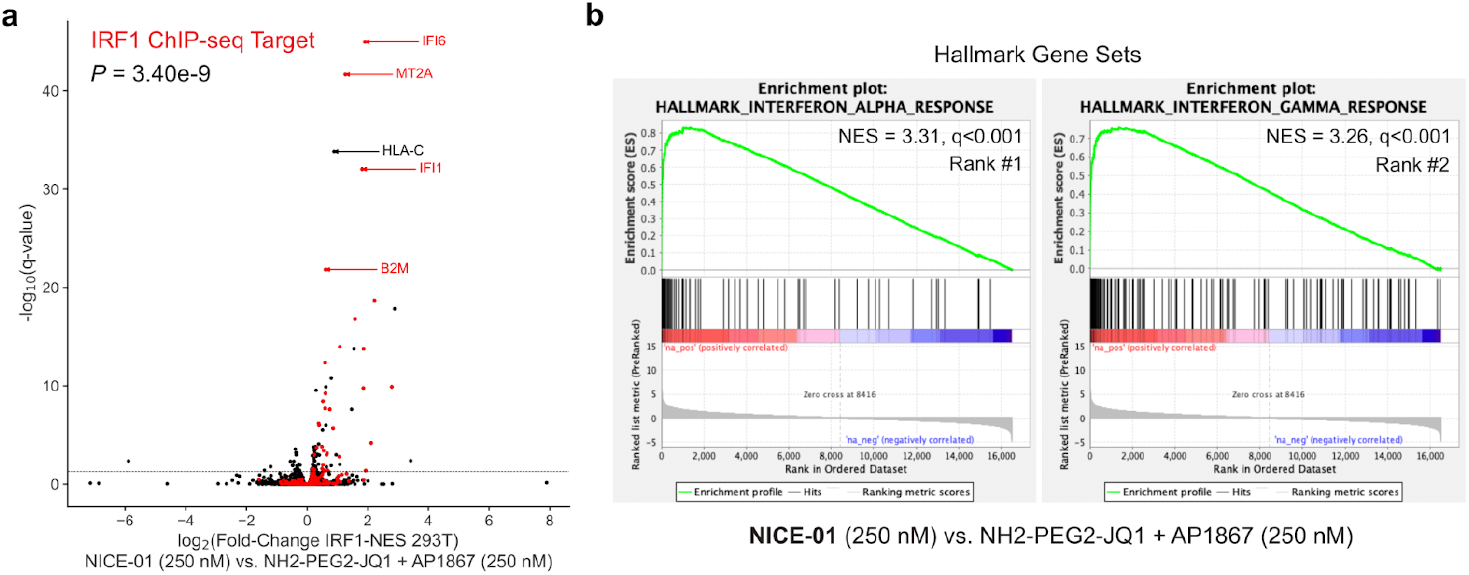
Import of cytosolic-localized transcription factor into the nucleus drives transcription. (A) 293T cells transfected with FKBP^F36V^-mEGFP-IRF1-NES and treated with **NICE-01** (250 nM) or JQ1-PEG2-NH2 + AP1867 (binders, 250 nM each), and subject to RNA-sequencing. Volcano plot showing upregulated genes following treatment with 250 nM **NICE-01** versus binders; genes in red are IRF1 ChIP-seq targets. *P*-value calculated from one sample t-test versus μ = 0. (B) Hallmark pre-ranked gene set enrichment analysis of cells treated with **NICE-01** versus binders. Top ranked positively enriched gene sets are shown.

## Discussion

The regulated localization of proteins into different cellular compartments is critical for proper cellular function. Here we use chemical-induced proximity between proteins that natively occupy different compartments to change localization of the targeted protein.

It was initially unclear whether inducing proximity between a nuclear protein and a cytosolic protein would cause the nuclear import of the cytosolic protein, the export of the nuclear protein or some equilibrium between the two. We had originally hypothesized for example that NPM1c would export BRD4 from the nucleus. In the nucleus, BRD4 makes contact with several large macromolecules such as the mediator complex, a 31-protein complex of proteins that collectively has 26 predicted annotated nuclear localization signals^15^. In this case, the transport equilibrium could heavily favor nuclear localization even when bound to an NES-containing protein like NPM1 due to a sort of NES-NLS voting mechanism. In principle, bifunctional compounds could transport nuclear proteins out of the nucleus if the mechanisms favoring export overcome those favoring import. Indeed, chemical-induced proximity with rapamycin has been used to deplete FRB-fused nuclear proteins from the nucleus through co-export with FKBP-fused ribosomal subunits, which display massive flux out of the nucleus during ribosome maturation^31^. Detailing bifunctional molecules that can localize proteins to the various compartments in the cell is an important area for future research.

We also observed that the stoichiometry of ternary complex components appears to impact nuclear import significantly. When relying on the relatively low levels of endogenous BRD4 for import of a highly expressed construct, we observed weaker nuclear enrichment of the target protein. Moreover, cells with high relative NPM1c expression compared to BRD4 were less likely to display NPM1c import. These findings lead us to believe compound-induced nuclear import could occur for two endogenously expressed proteins when their expression level is similar or when the nuclear-localized protein is in greater abundance. They also imply that if one uses a lineage-or cancer cell-specific nuclear protein as a “carrier”, one can restrict the target population of cells where the cargo is being imported. These stoichiometry restrictions likely apply to all induced-proximity systems requiring interactions over long time scales, such as molecular glues. For rapamycin, the abundance of the presenter (FKBP12, 2.9×10^6^ copies/cell in 293T; OpenCell^32^) is in far excess of the target (mTOR, 6.9×10^4^ copies/cell in 293T; OpenCell^32^). For systems only needing transient interactions, such as bifunctional molecules inducing post-translational modifications, this may be less relevant.

To our knowledge this is also the first observation of significant single-cell heterogeneity in the phenotypic response to bifunctional compounds. Stochastic variation in the abundance of proteins targeted by chemotherapeutics has previously been shown to underlie single-cell phenotypic heterogeneity^33^. This observation raises the possibility that similar heterogeneity may be widespread, and possibly therapeutically important for several emerging therapeutic classes of bifunctional compounds.

Because protein localization controls diverse biological phenomena, the ability to pharmacologically perturb localization may offer control over such biological phenomena. We believe this may have important therapeutic implications. For instance, NPM1c is an AML driver that induces dominant negative NPM1 cytosolic localization^18,20^. We attempted to correct this directly by showing it can be relocalized to the nucleus with a bifunctional small molecule. However, one may prefer relocalization to the nucleolus, where wild-type NPM1 normally resides; this may require a nucleolar carrier. Secondly, relocalization may not be sufficient to yield a phenotype, as evidenced by the lack of p53 co-import. In other cases, we hypothesize that future therapeutics may divert an oncogenic protein from its normal substrates. We showed that oncogenic PIK3CA mutant protein can be diverted from the plasma membrane where it normally interacts with phosphoinositols, into the nucleus. Future research should focus on the future applications and functional consequences of such relocalizations.

One notable observation during our experiments, especially with NPM1, was the change in the composition of cellular condensates. We noted relocalization of cytosolic and nucleolar NPM1c to BRD4 condensates. Therefore, chemical inducers of proximity may represent a modular strategy to alter condensate composition. BRD4 condensates are especially interesting due to their role in transcription^34^. We show that import of a cytosolic transcription factor with endogenous BET containing proteins can strongly induce transcription. This is consistent with results from previous studies using bifunctionals to recruit BRD4 to targeted sites^35–37^.

Overall, these data demonstrate that protein localization can be modulated in human cells by bifunctional molecules that bind proteins in different cellular compartments. The practical application of this approach, proved here using binders to FKBP^F36V^ and BRD4, will require the identification and optimization of protein-specific binding molecules. We anticipate that future bifunctional relocalizing molecules will have important applications in scientific discovery and human therapeutics.

## Acknowledgements

We dedicate this work to Peter J. Belshaw (1967-2021) whose doctoral studies both pioneered the powerful “bump-hole” concept and advanced the early phase “bifunctional compounds”. This work was supported by the NCI’s Cancer Target Discovery and Development (CTD2) Network (grant number U01CA217848, S.L.S), NCI grant R35CA197568 (M.M.), and an American Cancer Society Research Professorship (M.M.).

## Author Contributions

Conceptualization: W.J.G., A.S., Data Curation: W.J.G., A.S., V.M.S., Formal Analysis: W.J.G., A.S., Funding Acquisition: M.M., S.L.S., Investigation: W.J.G., A.S., V.M.S., Methodology: W.J.G., A.S., V.M.S., Project administration: W.J.G., A.S., Resources: A.C., M.M., S.L.S., Software: W.J.G., A.S., Supervision: A.C., M.M., S.L.S., Validation: W.J.G., A.S., Visualization: W.J.G., A.S., Writing – original draft: W.J.G., A.S., Writing – review & editing: W.J.G., A.S., V.M.S., A.C., M.M., S.L.S.

## Methods

### Cell Culture

293T and U2OS cells were obtained from ATCC. All cell lines were cultured in DMEM supplemented with 10% FBS, and 100 IU/mL of penicillin, 100 μg/mL of streptomycin at 37°C in 5% CO_2_.

### Transient Transfection

Cells were transfected using TransIT-LT1 following the manufacturer’s protocol.

### Microscopy

Cells were imaged using Operetta CLS High Content Analysis System (PerkinElmer) or Opera Phenix Plus High-Content Screening System (PerkinElmer) in 96-or 384-well plates at 37°C and 5% CO_2_. Cells were split into plates 24 hours prior to transfection. Cells were imaged 24-48 hours following transfection, with media being replaced immediately before imaging with FluoroBrite DMEM containing 10% FBS. Compounds were added to fresh FluoroBrite DMEM, vortexed for 30 seconds, and the media was then added to cells and they were re-imaged between 0 minutes to 24 hours after addition.

### RNA Sequencing

293T cells were transfected with FKBP^F36V^-mEGFP-IRF1-NES and treated with NICE-01 (250nM) or the combination of NH2-PEG2-JQ1 and AP1867 (250 nM each) for one day. Prior to RNA extraction, cells were washed once with cold PBS, TRIzol (Invitrogen) was added to cells, and RNA was extracted following the manufacturer’s protocol. RNA concentration was measured using a Qubit Fluorometer (ThermoFisher), and RNA integrity was assessed using an Agilent Bioanalyzer. RNA-sequencing libraries were prepared using the NEBNext Ultra II RNA Library Prep Kit (Illumina). Paired-end 150bp sequencing was done on a NovaSeq 6000 machine (Illumina).

### RNA-seq Analysis

STAR/RSEM was used to align paired end RNA-seq reads to the GENCODE v38 transcript reference. The raw counts were rounded, and DESeq2 was used to compare bifunctional versus binder treated replicates. We restricted to genes with baseMean ≥ 10 counts. A preranked GSEA was conducted using the DESeq2 t-statistic as the rank metric.

### Co-dependency Analysis

RNAi scores for NPM1 across DepMap were correlated with RNAi dependency scores for all other genes^38^. The resulting correlation coefficients were plotted versus q-values derived from Benjamini-Hochberg correction of Pearson correlation significance results.

### Comut Analysis

The OHSU acute myeloid leukemia cohort was queried on cBioPortal for *NPM1* and *TP53* mutations. A comut generated from cBioPortal is shown^39,40^.

### Microscopy Analysis

For kinetic analyses, ImageJ^41^ was used to quantify average cell mEGFP intensity (I_avg,cell_), cell area (A_cell_), average nuclear mEGFP intensity (I_avg,n_), nuclear area (A_n_), and background intensity (I_avg,b_). These parameters were used to calculate mEGFP cytosolic fraction: f_cyto_ = ((I_avg,cell_-I_avg,b_)A_cell_ -(I_avg,n_-I_avg,b_)A_n_) / (A_cell_(I_avg,cell_ - I_avg,b_)). CellProfiler^42^ was used for analysis of NPM1c transfected cells. Cells were identified as primary objects using a highlighted mCherry channel (which stains the entire cell due to detection of mCherry-BRD4 synthesized in the cytosol) with diameters of at least 15 pixels. The intensity and colocalization of non-enhanced (standard exposure) mCherry and mEGFP were computed in the identified cells. Integrated intensities and mCherry-mEGFP pixel Pearson correlation were used in downstream analysis.

## Supplementary Figures

**Figure S1.**
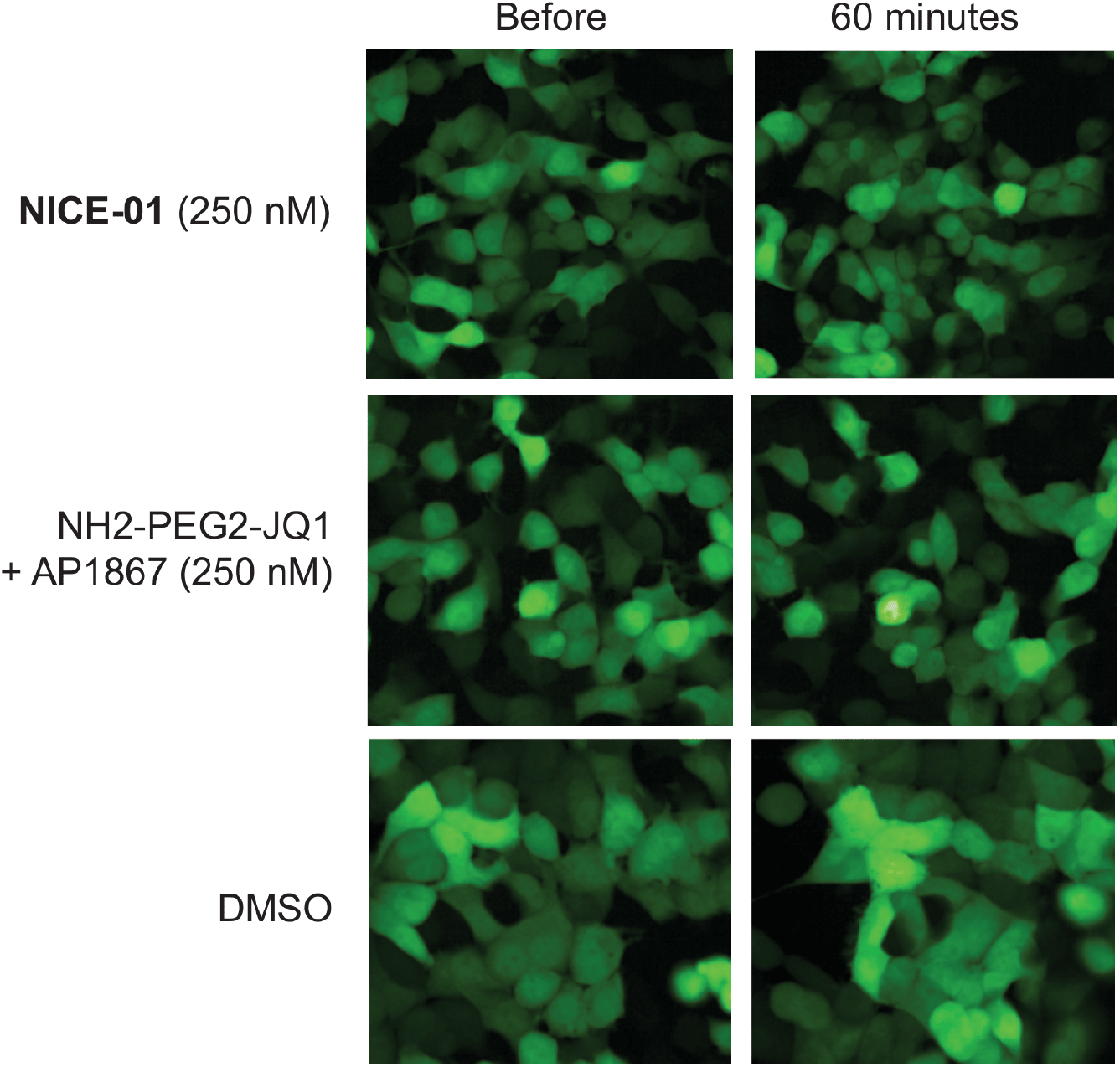
Import of GFP with endogenous BET-containing proteins. 293T cells transfected with FKBP^F36V^-mEGFP, treated with **NICE-01** (250 nM), NH2-PEG2-JQ1 + AP1867 (250 nM), or DMSO, and imaged for 60 minutes.

**Figure S2.**
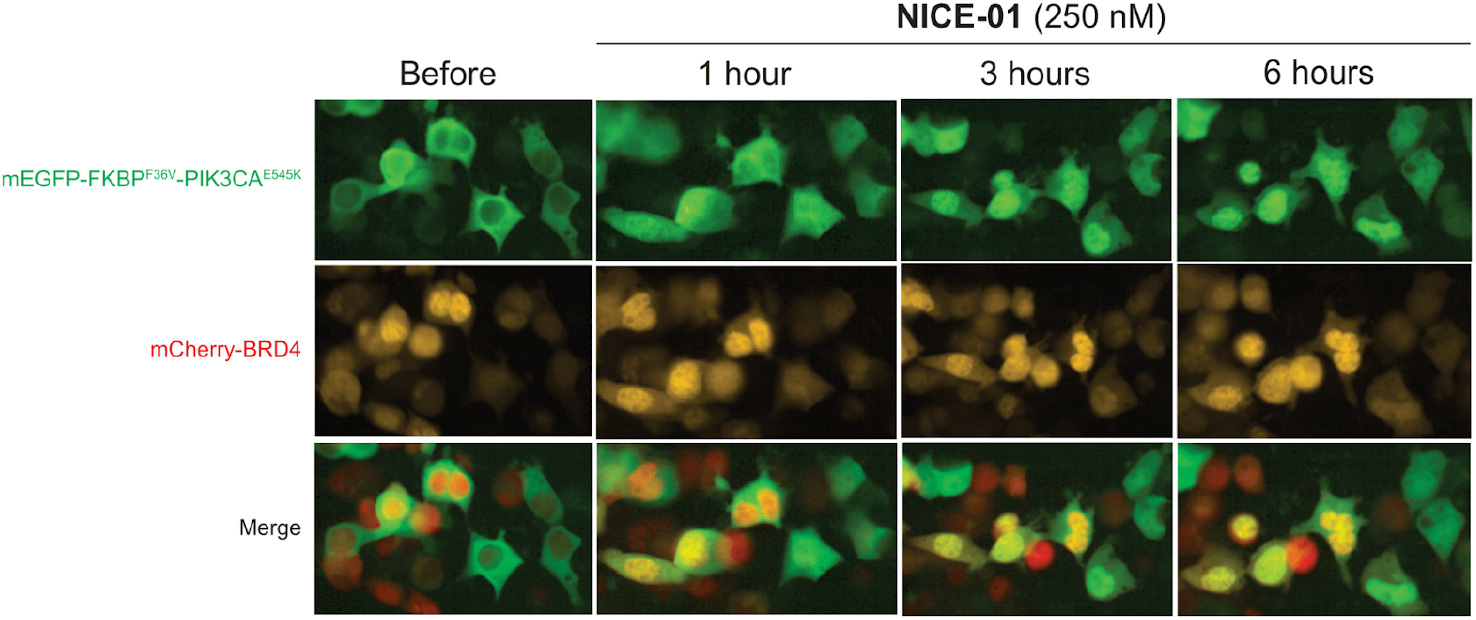
Import of a large cargo unable to diffuse across the nuclear pore. 293T cells co-transfected with mCherry-BRD4 and mEGFP-FKBP^F36V^-PIK3CA^E545K^, treated with **NICE-01** (250 nM) and imaged for 6 hours.

**Figure S3.**
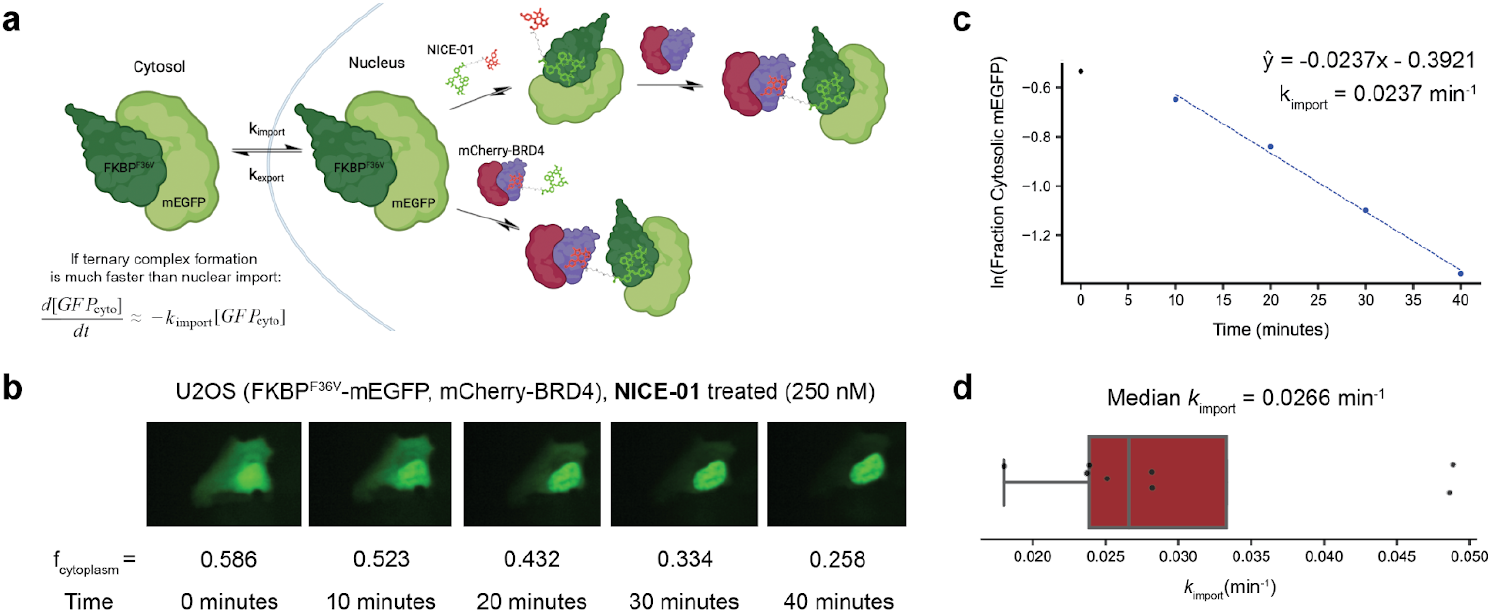
Kinetic parameters of nuclear import revealed by treatment with bifunctional **NICE-01**. (A) Schematic of mathematical model (see Supplementary Information). (B) U2OS cells co-transfected with mCherry-BRD4 and FKBP^F36V^-mEGFP, treated with **NICE-01** (250 nM), and imaged every 10 minutes for 40 minutes. Quantification of cytosolic mEGFP fraction at different timepoints. (C) ln(cytosolic mEGFP) versus time for representative cell shown in panel (**b**). (D) Boxplot of *k*_import_ for 8 cells with high coefficients of determination (R^2^ > 0.95). Each dot represents one cell.

**Figure S4.**
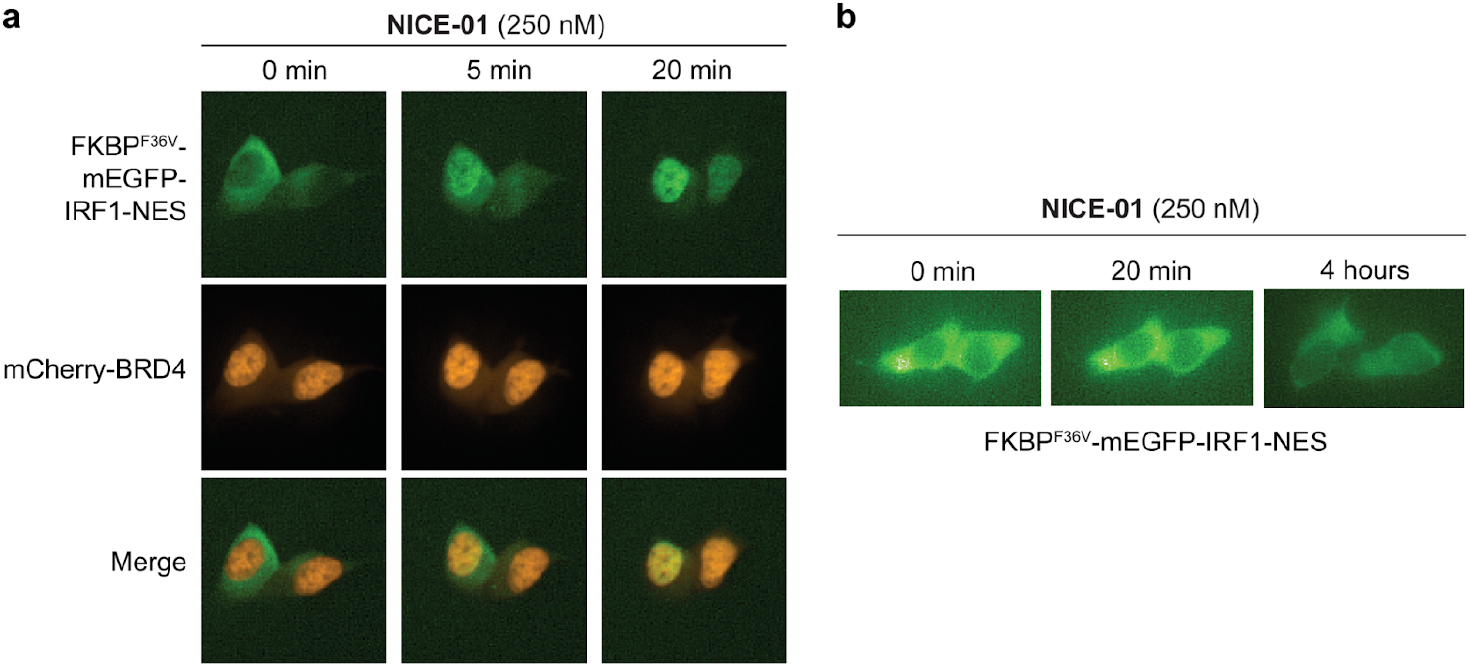
Assessment of IRF1-NES nuclear import with and without exogenous BRD4. (A) 293T cells co-transfected with FKBP^F36V^-mEGFP-IRF1-NES and mCherry-BRD4, treated with **NICE-01** (250 nM), and imaged for 20 minutes. (B) 293T cells transfected with FKBP^F36V^-mEGFP-IRF1-NES, treated with **NICE-01** (250 nM), and imaged for 4 hours.

## Supporting Information

### General Procedures

Commercial reagents were used for chemical synthesis without additional purification. Reactions were performed in round-bottom flasks with Teflon-coated magnetic stir bars. The progress of the reactions was tracked using ultra-performance liquid chromatography mass spectrometry. This analysis was conducted on a Waters ACQUITY UPLC I-Class 15 PLUS System with an ACQUITY SQ Detector 2. Nuclear magnetic resonance (NMR) spectra were obtained at the Broad Institute of MIT and Harvard using a Bruker AVANCE III HD 400 MHz spectrometer at room temperature (^1^H NMR, 400 MHz). Chemical shifts for 1H NMR are given in parts per million (ppm) and referenced to residual solvent signals. DMSO-d6 was acquired from Cambridge Isotope Laboratories, Inc. ^1^H NMR data is reported as follows: chemical shift value in ppm, multiplicity (s = singlet, d = doublet, t = triplet, dd = doublet of doublets, and m = multiplet), coupling constant value in Hz, and integration value.

### Organic Synthesis

#### (*S*)-N-(2-(2-(2-aminoethoxy)ethoxy)ethyl)-2-(4-(4-chlorophenyl)-2,3,9-trimethyl-6H-thieno[3,2-f][1,2,4]triazolo[4,3-a][1,4]diazepin-6-yl)acetamide (NH2-PEG2-JQ1)

To a solution of (+)-JQ1 (45.7 mg, 0.100 mmol, MedChemExpress) in dichloromethane (4 mL), trifluoroacetic acid (1 mL) was added and stirred overnight. After reaction completion, solvents were removed under reduced pressure. The residue was redissolved in DMF (1 mL). Amino-PEG2-amine (32 mg, 0.216 mmol), PyBOP (52 mg, 0.100 mmol), and DIPEA (35 μL) were added and stirred for 2 hours. The reaction was concentrated under reduced pressure and the product purified by reverse-phase HPLC (acetonitrile:water gradient up to 90%). After lyophilization of fractions, 31 mg of product was isolated (0.058 mmol, 58% yield). LC/MS of purified combined JQ1-PEG2-NH2 fractions is shown. ^1^H NMR spectra matched what was previously reported^43–45^. LRMS [M+H, C_25_H_32_ClN_6_O_3_S^+^] m/z theoretical 531.2, found 531.6.

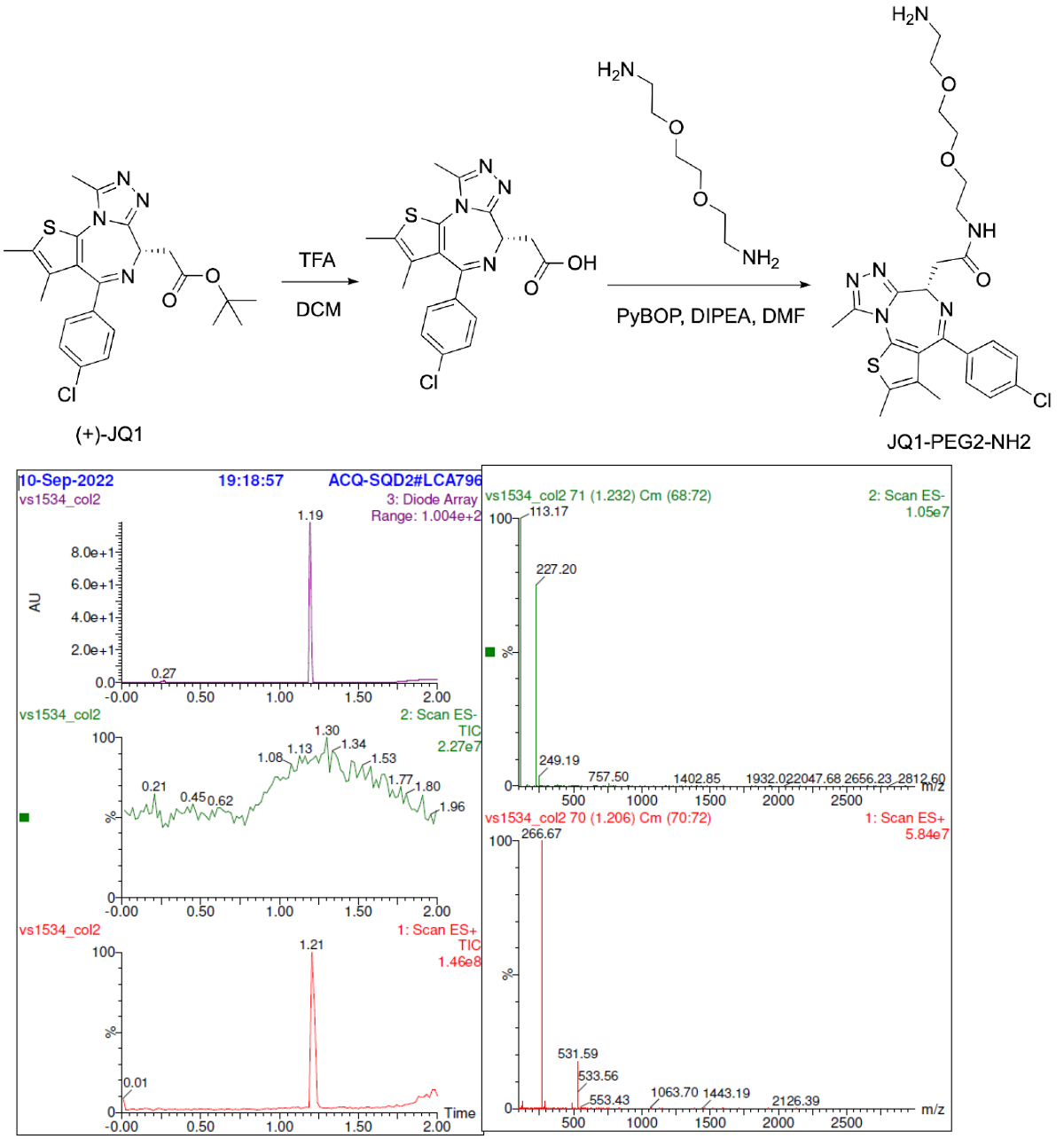

#### (*R*)-1-(3-((14-((*S*)-4-(4-chlorophenyl)-2,3,9-trimethyl-6H-thieno[3,2-f][1,2,4]triazolo[4,3-a][1,4]diazepin-6-yl)-2,13-dioxo-6,9-dioxa-3,12-diazatetradecyl)oxy)phenyl)-3-(3,4-dimethoxyphenyl)propyl (*S*)-1-((*S*)-2-(3,4,5-trimethoxyphenyl)butanoyl)piperidine-2-carboxylate (AP1867-PEG2-JQ1; AP-PEG2-JQ1; NICE-01)

To a solution of AP1867 (5 mg, 0.0072 mmol, MedChemExpress) in DMF (1 mL), JQ1-PEG2-NH2 (5 mg, 0.0094 mmol) and DIPEA (5 μL) were added and stirred for 30 minutes. After the reaction completion, the product was purified by reverse-phase HPLC (acetonitrile:water gradient up to 90%). After lyophilization of fractions, 5.0 mg of product was isolated (0.0041 mmol, 58% yield). LC/MS of purified combined AP-PEG2-JQ1 fractions is shown. ^1^H NMR (400 MHz, DMSO) δ 8.28 (t, *J* = 5.5 Hz, 1H), 8.12-8.05 (t, *J* = 5.2 Hz, 1H), 7.48 (d, *J* = 8.5 Hz, 2H), 7.42 (d, *J* = 8.2 Hz, 2H), 7.35-7.28 (minor rotamer) & 7.22-7.12 (major rotamer, m, 1H), 7.00-6.80 (m, 3H), 6.79-6.56 (m, 3H), 6.54 (s, 2H), 5.78 (minor rotamer) & 5.53 (major rotamer, dd, *J* = 8.4, 5.3 Hz, 1H), 5.28 (major rotamer, d, *J* = 4.2 Hz, 1H) & 5.05 (minor rotamer), 4.56-4.39 (m, 3H), 4.02 (d, *J* = 16.3 Hz, 2H), 3.87 (t, *J* = 7.2 Hz, 1H), 3.80-3.68 (m, 8H), 3.65-3.56 (m, 9H), 3.53 (s, 4H), 3.49-3.42 (m, 4H), 3.29-3.19 (m, 4H), 2.70-2.64 (m, 1H), 2.59 (s, 3H), 2.50-2.43 (m, 1H), 2.41 (s, 3H), 2.39-2.24 (m, 2H), 2.18 (d, *J* = 12.9 Hz, 2H), 2.00-1.86 (m, 2H), 1.62 (s, 3H), 1.57 (t, *J* = 7.1 Hz, 2H), 1.42-1.34 (m, 1H), 1.22-1.07 (m, 1H), 0.82 (major rotamer, t, *J* = 7.3 Hz, 3H) & 0.75 (minor rotamer). LRMS [M+H, C_63_H_77_ClN_7_O_13_S^+^] m/z theoretical 1206.5, found 1207.0.

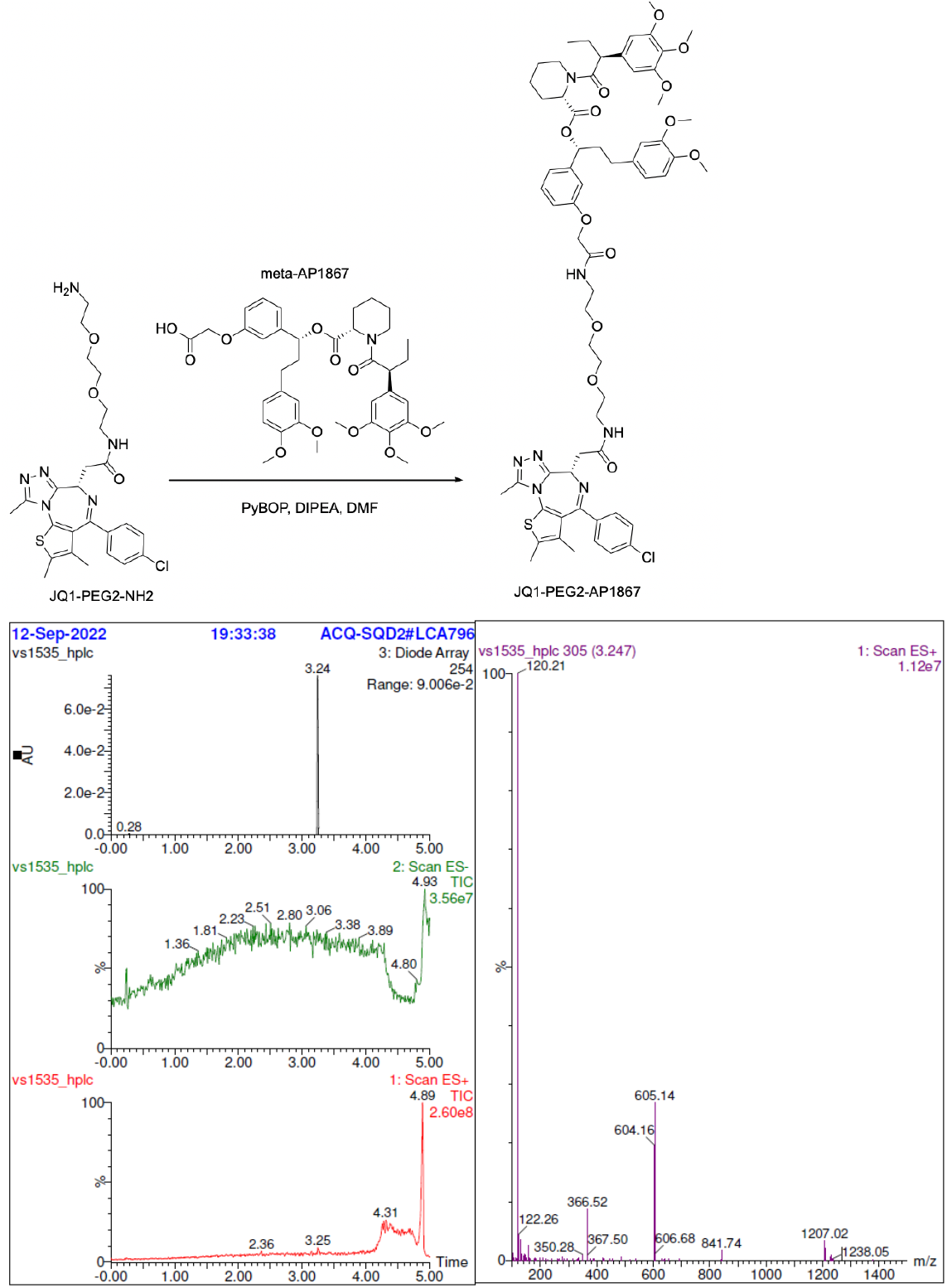

#### Kinetic Analysis

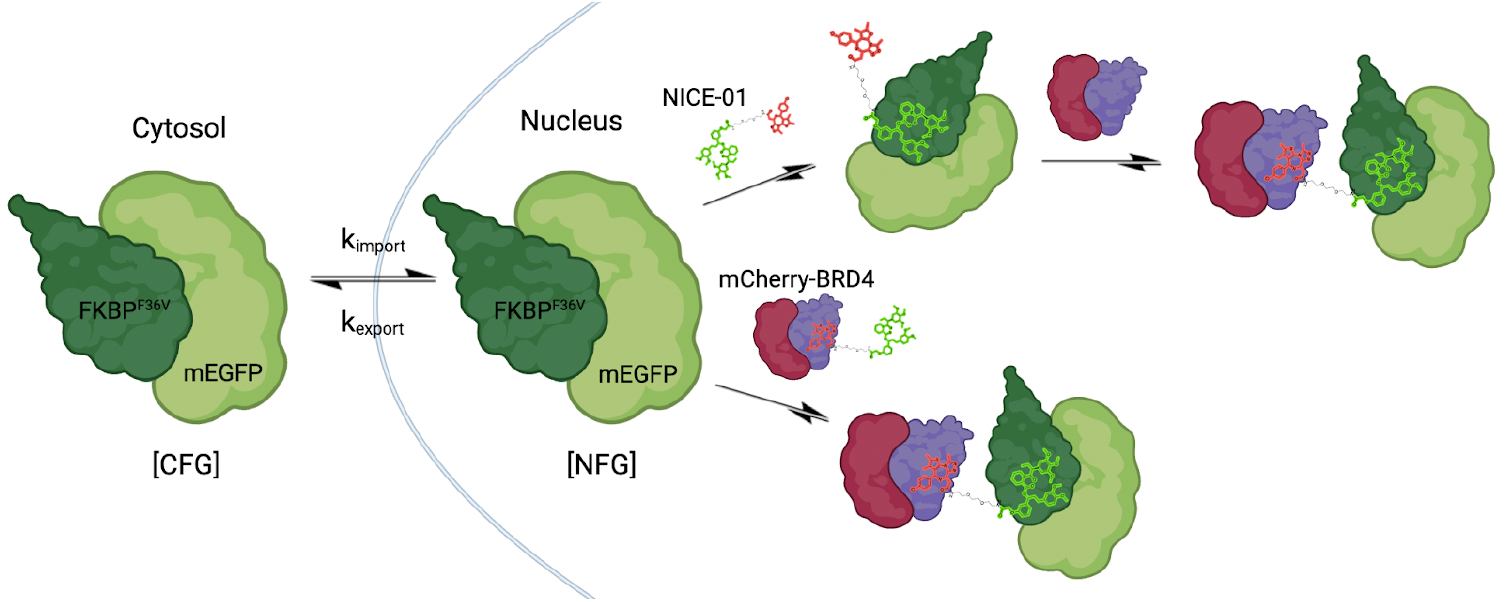

We ignore translation, degradation, and co-import due to the short time scale of experiments (40 minutes).

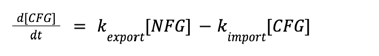

Prior to adding **NICE-01**, FKBP^F35V^-mEGFP is distributed approximately equally through the cytosol and nucleus (which indicates *k*_*import*_ =*k*_*export*_ =*k*_*diffudion*_, as expected to be the case with passive diffusion Import export diffusion across the nuclear pore). **NICE-01** causes the cytosolic fraction of mEGFP to decrease, which would not be possible in this system unless [*NFG*] decreases. If the rate of nuclear import is slow relative to the rate of ternary complex formation, we expect there to be a period of time where [*CFG*] ≫ [*NFG*]. During this time, if it exists, 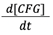 becomes first-order with respect to [CFG]:

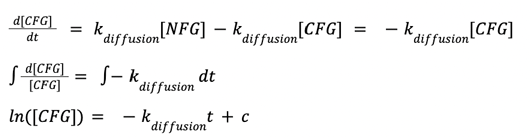

We define a constant: [*GFP*]_0_ = [*CFG*] +[*NFG*] +[*NMBFG*], this corresponds to the total concentration of GFP in the cell. This is assumed (and observed) to be constant throughout the experiment.

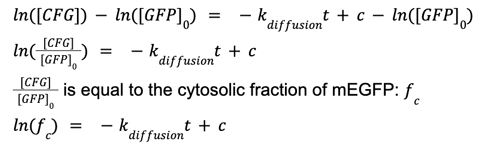

Therefore, the observation of a linear relationship on the *ln*(*f*_*c*_) versus time curve that arise after compound addition in most cells, suggests nuclear import is rate-limiting. The slope of this linear portion is equal to the rate constant for diffusion across the nuclear pore for FKBP^F36V^-mEGFP.

#### Schematics of Plasmids

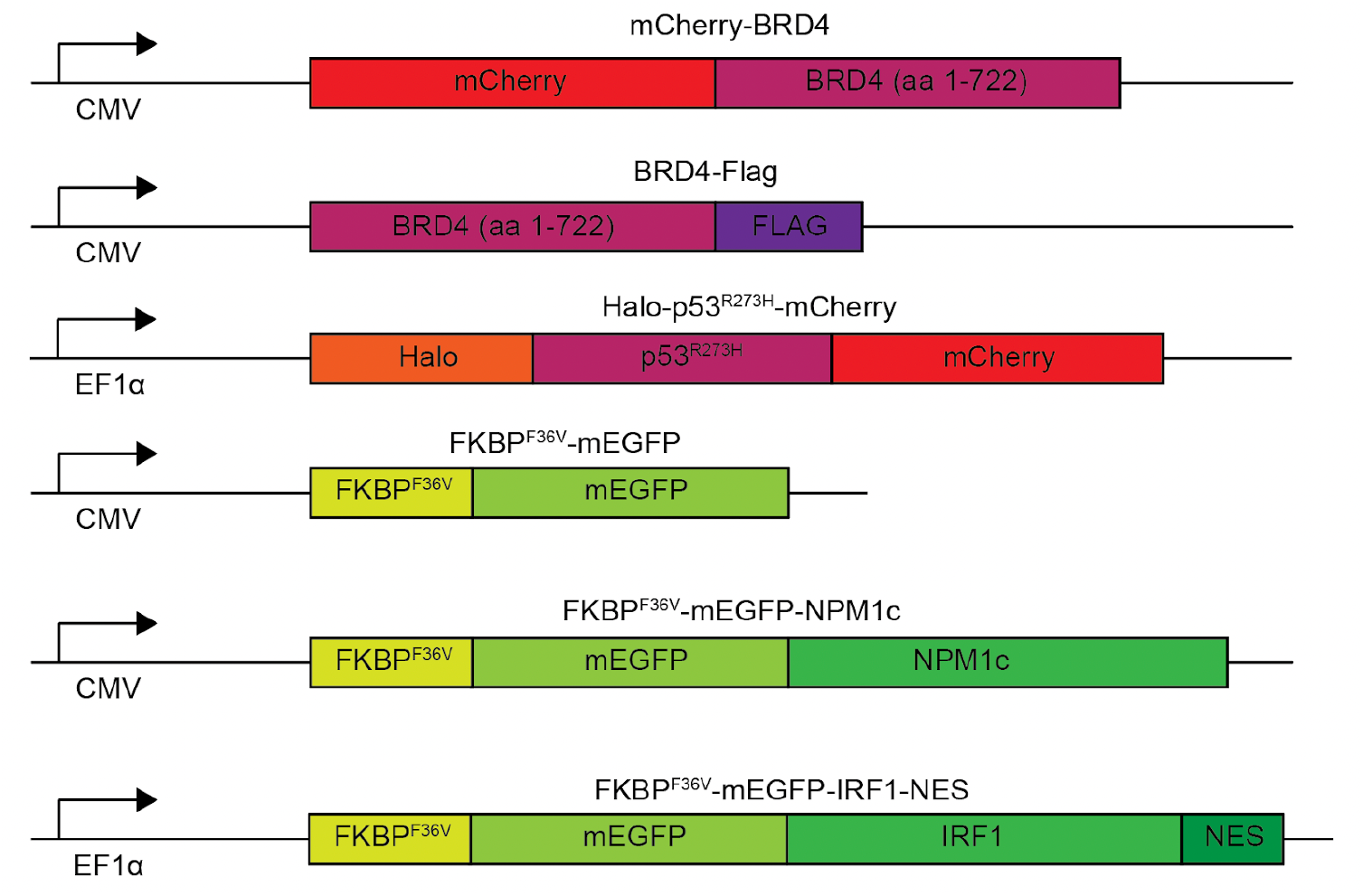

## References

1. Kalderon, D.; Roberts, B. L.; Richardson, W. D.; Smith, A. E. A Short Amino Acid Sequence Able to Specify Nuclear Location. Cell 1984, 39 (3 Pt 2), 499–509.

2. Wang, R.; Brattain, M. G. The Maximal Size of Protein to Diffuse through the Nuclear Pore Is Larger than 60kDa. FEBS Lett. 2007, 581 (17), 3164–3170.

3. Banani, S. F.; Lee, H. O.; Hyman, A. A.; Rosen, M. K. Biomolecular Condensates: Organizers of Cellular Biochemistry. Nat. Rev. Mol. Cell Biol. 2017, 18 (5), 285–298.

4. Jiang, X.; Wang, X. Cytochrome C-Mediated Apoptosis. Annu. Rev. Biochem. 2004, 73, 87–106.

5. Ivanov, P.; Kedersha, N.; Anderson, P. Stress Granules and Processing Bodies in Translational Control. Cold Spring Harb. Perspect. Biol. 2019, 11 (5).

6. Van Der Heide, L. P.; Hoekman, M. F. M.; Smidt, M. P. The Ins and Outs of FoxO Shuttling: Mechanisms of FoxO Translocation and Transcriptional Regulation. Biochem. J 2004, 380 (Pt 2), 297–309.

7. Meyer, T.; Vinkemeier, U. STAT Nuclear Translocation: Potential for Pharmacological Intervention. Expert Opin. Ther. Targets 2007, 11 (10), 1355–1365.

8. Sever, R.; Glass, C. K. Signaling by Nuclear Receptors. Cold Spring Harb. Perspect. Biol. 2013, 5 (3), a016709.

9. Belshaw, P. J.; Ho, S. N.; Crabtree, G. R.; Schreiber, S. L. Controlling Protein Association and Subcellular Localization with a Synthetic Ligand That Induces Heterodimerization of Proteins. Proc. Natl. Acad. Sci. U. S. A. 1996, 93 (10), 4604–4607.

10. Winter, G. E.; Buckley, D. L.; Paulk, J.; Roberts, J. M.; Souza, A.; Dhe-Paganon, S.; Bradner, J. E. Phthalimide Conjugation as a Strategy for in Vivo Target Protein Degradation. Science 2015, 348 (6241), 1376–1381.

11. Koide, K.; Finkelstein, J. M.; Ball, Z.; Verdine, G. L. A Synthetic Library of Cell-Permeable Molecules. J. Am. Chem. Soc. 2001, 123 (3), 398–408.

12. Zorba, A. et al. Delineating the Role of Cooperativity in the Design of Potent PROTACs for BTK. Proc. Natl. Acad. Sci. U. S. A. 2018, 115 (31), E7285–E7292.

13. Raina, K. et al. Regulated Induced Proximity Targeting Chimeras (RIPTACs): A Novel Heterobifunctional Small Molecule Therapeutic Strategy for Killing Cancer Cells Selectively. bioRxiv, 2023, https://doi.org/10.1101/2023.01.01.522436.

14. Kolos, J. M.; Voll, A. M.; Bauder, M.; Hausch, F. FKBP Ligands-Where We Are and Where to Go? Front. Pharmacol. 2018, 9, 1425.

15. Tessier, T. M.; MacNeil, K. M.; Mymryk, J. S. Piggybacking on Classical Import and Other Non-Classical Mechanisms of Nuclear Import Appear Highly Prevalent within the Human Proteome. Biology 2020, 9 (8).

16. Mohr, D.; Frey, S.; Fischer, T.; Güttler, T.; Görlich, D. Characterisation of the Passive Permeability Barrier of Nuclear Pore Complexes. EMBO J. 2009, 28 (17), 2541–2553.

17. Falini, B.; Brunetti, L.; Sportoletti, P.; Martelli, M. P. NPM1-Mutated Acute Myeloid Leukemia: From Bench to Bedside. Blood 2020, 136 (15), 1707–1721.

18. Cancer Genome Atlas Research Network. Genomic and Epigenomic Landscapes of Adult de Novo Acute Myeloid Leukemia. N. Engl. J. Med. 2013, 368 (22), 2059–2074.

19. Falini, B. et al. Both Carboxy-Terminus NES Motif and Mutated Tryptophan(s) Are Crucial for Aberrant Nuclear Export of Nucleophosmin Leukemic Mutants in NPMc+ AML. Blood 2006, 107 (11), 4514–4523.

20. Bolli, N. et al. Born to Be Exported: COOH-Terminal Nuclear Export Signals of Different Strength Ensure Cytoplasmic Accumulation of Nucleophosmin Leukemic Mutants. Cancer Res. 2007, 67 (13), 6230–6237.

21. Brunetti, L. et al. Mutant NPM1 Maintains the Leukemic State through HOX Expression. Cancer Cell 2018, 34 (3), 499–512.e9.

22. Holoubek, A.; Strachotová, D.; Otevřelová, P.; Röselová, P.; Heřman, P.; Brodská, B. AML-Related NPM Mutations Drive p53 Delocalization into the Cytoplasm with Possible Impact on p53-Dependent Stress Response. Cancers 2021, 13 (13).

23. Barretina, J. et al. The Cancer Cell Line Encyclopedia Enables Predictive Modelling of Anticancer Drug Sensitivity. Nature 2012, 483 (7391), 603–607.

24. Tyner, J. W. et al. Functional Genomic Landscape of Acute Myeloid Leukaemia. Nature 2018, 562 (7728), 526–531.

25. Ferrigno, P.; Silver, P. A. Regulated Nuclear Localization of Stress-Responsive Factors: How the Nuclear Trafficking of Protein Kinases and Transcription Factors Contributes to Cell Survival. Oncogene 1999, 18 (45), 6129–6134.

26. Lin, R.; Heylbroeck, C.; Pitha, P. M.; Hiscott, J. Virus-Dependent Phosphorylation of the IRF-3 Transcription Factor Regulates Nuclear Translocation, Transactivation Potential, and Proteasome-Mediated Degradation. Mol. Cell. Biol. 1998, 18 (5), 2986–2996.

27. Forero, A. et al. Differential Activation of the Transcription Factor IRF1 Underlies the Distinct Immune Responses Elicited by Type I and Type III Interferons. Immunity 2019, 51 (3), 451–464.e6.

28. Tanaka, N.; Kawakami, T.; Taniguchi, T. Recognition DNA Sequences of Interferon Regulatory Factor 1 (IRF-1) and IRF-2, Regulators of Cell Growth and the Interferon System. Mol. Cell. Biol. 1993, 13 (8), 4531–4538.

29. Rettino, A.; Clarke, N. M. Genome-Wide Identification of IRF1 Binding Sites Reveals Extensive Occupancy at Cell Death Associated Genes. J Carcinog Mutagen 2013.

30. Subramanian, A. et al. Gene Set Enrichment Analysis: A Knowledge-Based Approach for Interpreting Genome-Wide Expression Profiles. Proc. Natl. Acad. Sci. U. S. A. 2005, 102 (43), 15545–15550.

31. Haruki, H.; Nishikawa, J.; Laemmli, U. K. The Anchor-Away Technique: Rapid, Conditional Establishment of Yeast Mutant Phenotypes. Mol. Cell 2008, 31 (6), 925–932.

32. Cho et al. OpenCell: Endogenous Tagging for the Cartography of Human Cellular Organization. Science 2022, 375 (6585), eabi6983.

33. Spencer, S. L.; Gaudet, S.; Albeck, J. G.; Burke, J. M.; Sorger, P. K. Non-Genetic Origins of Cell-to-Cell Variability in TRAIL-Induced Apoptosis. Nature 2009, 459 (7245), 428–432.

34. Sabari, B. R. et al. Coactivator Condensation at Super-Enhancers Links Phase Separation and Gene Control. Science 2018, 361 (6400).

35. Erwin, G. S. et al. Synthetic Transcription Elongation Factors License Transcription across Repressive Chromatin. Science 2017, 358 (6370), 1617–1622.

36. Gourisankar S. et al. Rewiring Cancer Drivers to Activate Apoptosis. bioRxiv. https://doi.org/10.1101/2022.12.04.517548.

37. Chiarella, A. M. et al. Dose-Dependent Activation of Gene Expression Is Achieved Using CRISPR and Small Molecules That Recruit Endogenous Chromatin Machinery. Nat. Biotechnol. 2020, 38 (1), 50–55.

38. Tsherniak, A. et al. Defining a Cancer Dependency Map. Cell 2017, 170 (3), 564–576.e16.

39. Cerami, E. et al. The cBio Cancer Genomics Portal: An Open Platform for Exploring Multidimensional Cancer Genomics Data. Cancer Discov. 2012, 2 (5), 401–404.

40. Gao, J. et al. Integrative Analysis of Complex Cancer Genomics and Clinical Profiles Using the cBioPortal. Sci. Signal. 2013, 6 (269), 1.

41. Schneider, C. A.; Rasband, W. S.; Eliceiri, K. W. NIH Image to ImageJ: 25 Years of Image Analysis. Nat. Methods 2012, 9 (7), 671–675.

42. Stirling, D. R.; Swain-Bowden, M. J.; Lucas, A. M.; Carpenter, A. E.; Cimini, B. A.; Goodman, A. CellProfiler 4: Improvements in Speed, Utility and Usability. BMC Bioinformatics 2021, 22 (1), 433.

## Supporting References

43. Ren, C.; Zhang, G.; Han, F.; Fu, S.; Cao, Y.; Zhang, F.; Zhang, Q.; Meslamani, J.; Xu, Y.; Ji, D.; Cao, L.; Zhou, Q.; Cheung, K.-L.; Sharma, R.; Babault, N.; Yi, Z.; Zhang, W.; Walsh, M. J.; Zeng, L.; Zhou, M.-M. Spatially Constrained Tandem Bromodomain Inhibition Bolsters Sustained Repression of BRD4 Transcriptional Activity for TNBC Cell Growth. Proc. Natl. Acad. Sci. U. S. A. 2018, 115 (31), 7949–7954.

44. Richardson, P. L.; Marin, V. L.; Koeniger, S. L.; Baranczak, A.; Wilsbacher, J. L.; Kovar, P. J.; Bacon-Trusk, P. E.; Cheng, M.; Hopkins, T. A.; Haman, S. T.; Vasudevan, A. Controlling Cellular Distribution of Drugs with Permeability Modifying Moieties. Medchemcomm 2019, 10 (6), 974–984.

45. Henning, N. J.; Manford, A. G.; Spradlin, J. N.; Brittain, S. M.; Zhang, E.; McKenna, J. M.; Tallarico, J. A.; Schirle, M.; Rape, M.; Nomura, D. K. Discovery of a Covalent FEM1B Recruiter for Targeted Protein Deg-radation Applications. J. Am. Chem. Soc. 2022, 144 (2), 701–708.

